# Targeting DOT1L and EZH2 synergizes in breaking the germinal center identity of Diffuse Large B Cell Lymphoma

**DOI:** 10.1101/2024.06.05.597556

**Authors:** Camiel Göbel, Rachele Niccolai, Marnix H.P. de Groot, Jayashree Jayachandran, Joleen Traets, Ben Morris, Ji-Ying Song, Leyla Azarang, Eirini Kasa, Daniel de Groot, Maaike Kreft, Marie José Kersten, Muhammad A. Aslam, Fred van Leeuwen, Heinz Jacobs

## Abstract

Differentiation of antigen-activated B cells into pro-proliferative germinal center (GC) B cells depends on the activity of the transcription factor MYC and the epigenetic writers DOT1L and EZH2. GCB-like Diffuse Large B Cell Lymphomas (GCB-DLBCLs) arise from GC B cells and closely resemble their cell of origin. Given the dependency of GC B cells on DOT1L and EZH2, we investigated the role of these epigenetic regulators in GCB-DLBCL cell lines and observed that GCB-DLBCLs synergistically depend on the combined activity of DOT1L and EZH2. Mechanistically, inhibiting both enzymes led to enhanced derepression of PRC2 target genes compared to EZH2 single treatment, along with the suppression of MYC target genes. The sum of all these alterations results in a ‘cell-identity crisis,’ wherein GCB-DLBCLs lose their pro-proliferative GC identity and partially undergo plasma cell differentiation, a state associated with poor survival. In support of this model, combined epi-drugging of DOT1L and EZH2 prohibited the outgrowth of human GCB-DLBCL xenografts *in vivo*. We conclude that the malignant behavior of GCB-DLBCLs strictly depends on DOT1L and EZH2 and that combined targeting of both epigenetic writers may provide an alternative differentiation-based treatment modality for GCB-DLBCL.

**Key points:**

1. MYC-driven GCB-DLBCL depends on DOT1L and EZH2 and combined targeting provides an alternative differentiation-based therapy.
2. DOT1L and EZH2 cooperatively repress PRC2 target genes which is essential for acquiring a plasma cell-like state in GC B cell-like DLBCL.

## Introduction

Upon antigen encounter, naïve resting B cells get activated and differentiate into pro-proliferative, pro-mutagenic germinal center B cells in a T-cell dependent manner and become part of a specialized micro-environment, known as the Germinal Center (GC)^1–4^. Survivors of the GC reaction either differentiate into memory B cells to establish long-term immunological memory or plasma cells (PCs) ^5^. The pro-mutagenic activities in GC B cells are closely related to recurrent oncogenic chromosomal translocations and amplifications of proto-oncogenes like *MYC* and *BCL2*, rendering GC B cells prone to oncogenic transformation ^6^. Despite the effectiveness of rituximab plus cyclophosphamide, doxorubicin, vincristine, and prednisone (R-CHOP) chemoimmunotherapy, the standard of care treatment for DLBCL, approximately 40 percent of patients experience refractory disease or relapse ^7^. Notably, patients with *MYC* translocated DLBCL (5% to 14% of the cases) have a poor prognosis with R-CHOP therapy, highlighting the clinical need for alternative treatment approaches ^8,9^.

GC B cell function and behavior are closely associated with defined epigenetic changes mediated by the H3K27 methyltransferase Enhancer of Zeste Homolog 2 (EZH2), the catalytic component of the Polycomb Repressive Complex 2 (PRC2) ^10–12^. The reliance of GC B cells on EZH2 is maintained in GC B cell-derived diffuse large B cell lymphoma (GCB-DLBCL), and gain-of-function EZH2 mutations drive the malignant transformation, support cell survival, and repress differentiation ^13–15^. EZH2 inhibitors such as tazemetostat have already been evaluated in a phase II clinical trial with an objective response rate of 69% and 35% in *EZH2 gain-of-function* and *EZH2 wild-type* follicular lymphoma patients, respectively ^16^. Independently, the H3K79 methyltransferase DOT1L was identified as a master regulator of the GC reaction ^17,18^. Loss of *Dot1L* in the mouse B cell lineage prohibited the formation of GCs and instead supported premature acquisition of a dysfunctional plasma cell-like state incompatible with survival. DOT1L gained wide attention as a dependency and specific drug target in MLL-rearranged leukemia, although the clinical efficacy of the DOT1L inhibitor EPZ-5676 as monotherapy was modest in a phase I trial ^19^.

Given the fact that GC B cells heavily depend on the activity of DOT1L and EZH2 to establish and maintain their pro-proliferative identity, we hypothesized that, based on a common cell of origin, GCB-DLBCL also rely on the activity of these key epigenetic writers. Our data suggest that the majority of GCB-DLBCLs is hyperresponsive to the co-inhibition of DOT1L and EZH2, offering an alternative combinatorial epi-drug and differentiation-based treatment modality for patients with DLBCL.

## Methods

### Cell lines and reagents

GCB-DLBCL cell lines were cultured in Iscove’s Modified Dulbecco Medium (IMDM) supplemented with 8% fetal calf serum, 1% GlutaMAX, and penicillin/streptomycin. Cells were maintained at 37°C and 5% CO_2_. EPZ-5676 (cat. No. HY-15593), GSK343 (cat. No. HY-13500), tazemetostat (cat. No. HY-13803), and DOT1L-IN-5 (cat. No. HY-135128) were obtained from MedChemExpress.

### Dose response curves and synergy experiments

A semi-automated seeding method was used to determine the dose-response curves and synergistic potential. 5×10^4 cells/mL were reseeded with fresh inhibitors and medium every four days until day 12 using a robotics protocol (Hamilton CO-RE96 Platform) and treated with EPZ-5676 (DOT1Li) and/or GSK343 (EZH2i) using the drug dispenser (HP Tecan digital dispenser). Cells were stained with DAPI to measure the viable cells by flow cytometry. The combination index (CI) was calculated using CompuSyn software. CI < 0.7 = synergism, 0.7 < CI < 1.1 = additive, and CI > 1.1 = antagonism ^20^.

### Flow cytometry

For intracellular staining of H3K27me3 (Cell Signaling, cat. No. 12158S), EZH2 (Cell Signaling, cat. No. 30233S), and IgM (BioLegend, cat. No. 314506), cells were fixed and permeabilized using the Transcription Factor Buffer set (BD Biosciences, cat. No. 562574) according to manufacturer’s protocol. H3K79me2 (Cell Signaling, cat. No. CST 5427S) staining was performed as previously described by Aslam et al ^17^. Flow cytometry was performed using the LSR Fortessa (BD Biosciences), and data were analyzed with FlowJo software (Tree Star Inc.).

### Mitochondrial membrane potential and apoptosis

GCB-DLBCL cell lines were treated with EPZ-5676 and GSK343 for six days. At day 3, 5×10^4 cells/ml were reseeded with fresh medium and inhibitors, including 25µM z-VAD-fmk (SelleckChem, cat. No. S7023). At day 6, cells were incubated with 1:125 TMRE (tetramethylrhodamine ethyl ester) (10µM) (Invitrogen™, cat. No. T669) for 30 minutes at 37°C in PBC + 1% FCS. Cells were washed with warm PBS + 1%FCS and incubated with 1:50 DAPI (70µg/ml, Sigma-Aldrich, cat. No. D9542) and 1:50 Annexin V-APC (Biolegend, cat. No. 640920) in 1x binding buffer (BD Biosciences, cat. No. 556454) for 15 minutes at room temperature and analyzed by flow cytometry.

### RNA sequencing and analysis

Cells were lysed in RLT buffer (Qiagen). RNA-seq libraries were prepared using the xGen polyA+ stranded RNA library kit, according to manufacturer’s protocol. Paired-end 54bp sequencing was performed on the Illumina NovaSeq 6000, on a SP flowcell. FASTQ files were produced by BCL Convert using demultiplexing. Sample distribution was based on indexing using IDT UDI indexes. Seqpurge was used for adapter trimming. RNA-seq data were aligned with STAR against reference genome GRCh38, and reads were counted and annotated per gene using GenSum (https://www.github.com/NKI-GCF/gensum) against Ensembl GRCh38.107. DESeq2 package was used to normalize read counts and to obtain the differential gene expression of RNA-seq samples.

### Gene signature analysis

To obtain gene signatures for naïve B cells (NBC), centroblasts (CB), plasmablasts (PB), memory B cells (MBC), and bone marrow plasma cells (BMPC), we took advantage of the ‘human B cells to plasma cells GCRMA’ dataset from GenomicScape (http://www.genomicscape.com/) ^21,22^. We obtained the top 500 genes of each B cell subset that were significantly upregulated compared to the other subsets (FDR < 0.05; fold change > 2). Gene Set Variation Analysis was performed using the GSVA R package (version 3.18) ^23^.

### ChIP sequencing, processing, and analysis

H3K27me3 ChIP-Seq sample and library preparation of Oci-Ly7 and Oci-Ly8 cell lines were performed as described by Aslam et al ^17^. Preprocessing of raw paired-end FASTA files was performed using the nf-core ATAC pipeline (version 2.0, https://nf-co.re/atacseq/2.0/) default settings ^24^. Specifically, Trim Galore! (version 0.6.7) was performed for adapter trimming before alignment. Reads were aligned against reference genome GRCh38 (Ensembl v109) with BWA (version 0.7.17). Broad/narrow peaks were called with MACS2 (version 2.2.7.1). Quality controls include FastQC (version 0.11.9), Samtools (version 1.15.1), Picard (version 2.27.4), and DeepTools (version 3.5.1). For downstream analysis, we used the merged BigWig file of filtered alignments across three biological replicates and normalized by scaling to 1 million mapped reads.

### In vivo tumor control of Oci-Ly7 xenografts

Six to eight-week-old NOD-Scid IL2Rgnull (Jackson Laboratory) mice, 50% males/females, were injected subcutaneously in the right flank with 1×10^7 Oci-Ly7 cells in 50% Matrigel (Corning) in PBS. Treatment started when tumors reached a volume of 80-100 mm^3^. Mice were randomized to treatment with vehicle, DOT1L-IN-5 (75mg/kg in Kolliphor HS15 by IP injection), tazemetostat (300mg/kg in 0.5% HPMC+ 0.1% Tween-80 in water by oral gavage), or the combination. Compounds were administrated twice daily with seven hours in between treatments. Mice were sacrificed when tumors reached 1500mm^3^ or on day 24, whichever came first.

Additional procedures and methods are detailed in the supplemental Methods.

## Results

### DOT1L inhibition negatively affects cell proliferation in a subset of GCB-DLCBL cell lines

Mouse GC B cells depend on *Dot1L* and *Ezh2* for their cellular identity ^17,18^. Based on cell of origin classification, we rationalized that human GCB-DLBCLs also rely on DOT1L to maintain their pro-proliferative identity. To evaluate the responsiveness of GCB-DLBCL to DOT1L inhibitor EPZ-5676 (Pinometostat; DOT1Li), we treated a panel of nine human GCB-DLBCL cell lines for 12 days with DOT1Li. Based on the dose-response curves, a clear distinction could be made between DOT1Li-responsive cell lines and -non responsive cell lines wherein EC_50_ values <150nM were associated with DOT1Li responsiveness **(Figure 1A)**. DOT1Li sensitivity did not correlate with the major genetic alterations reported for these cell lines, indicating that responsiveness to DOT1Li is unlikely predictable based on recurrent oncogenic alterations found in DLBCLs **(supplemental Table 1)**.

**Figure 1.**
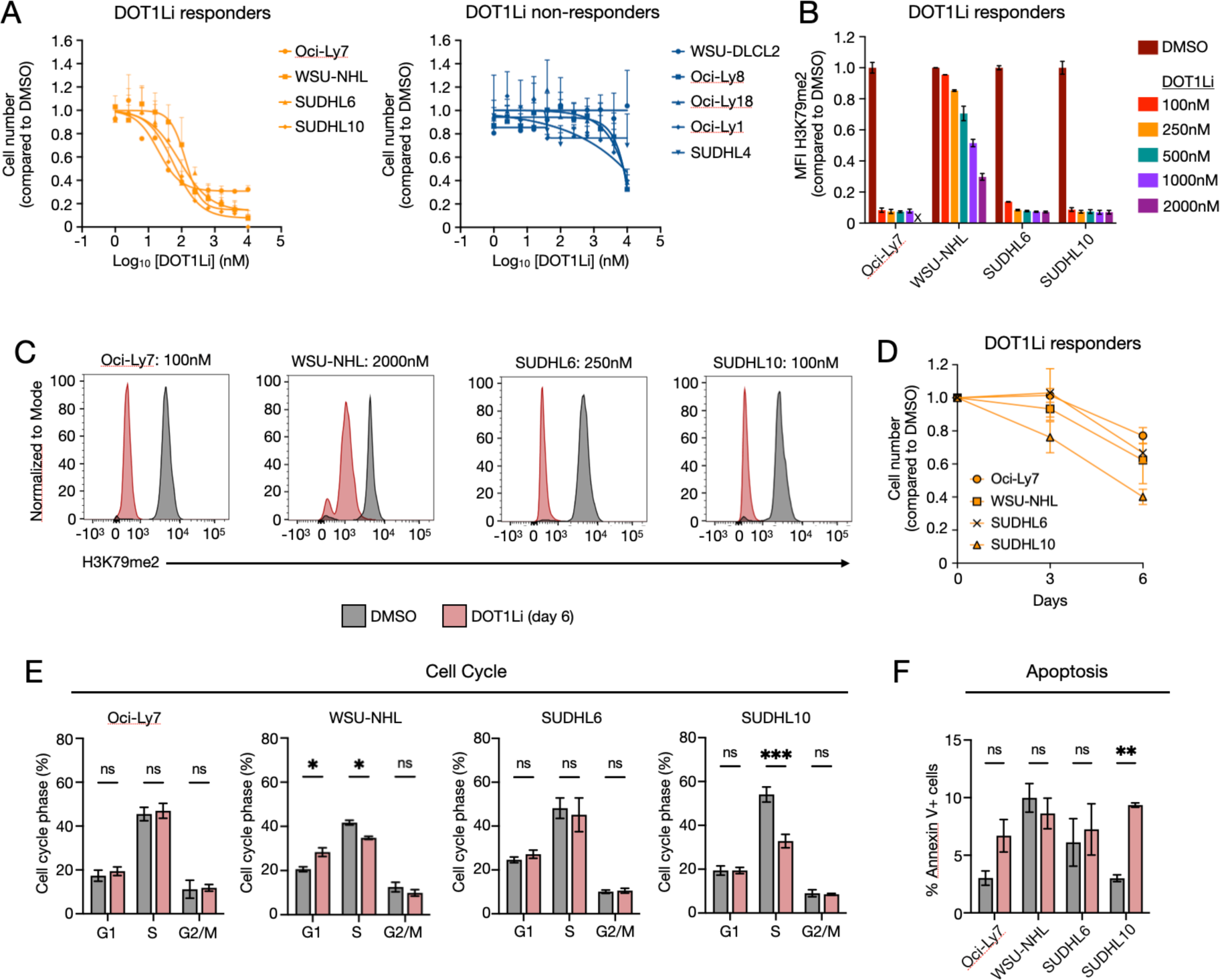
DOT1L inhibition negatively affects proliferation in a subset of GCB-DLBCL. (A) Dose-response curves of sensitive and non-responsive GCB-DLBCL cell lines towards DOT1L inhibition after 12 days of treatment. (B) Median fluorescence intensity (MFI) of H3K79me2 after six days of EPZ-5676 treatment in DOT1Li-sensitive GCB-DLBCL cell lines. Results represent two biological replicates. (C) Representative histograms showing the lowest dose required for viable cells to maximally reduce H3K79me2 after DOT1Li treatment. (D) Proliferation curves of DOT1Li-sensitive GCB-DLBCL cell lines upon DOT1Li treatment. Results represent two biological replicates. (E) Bar graphs showing the impact of DOT1L inhibition on cell cycle phases G1, S, and G2/M. *P*-values were calculated using two-way ANOVA. (F) Effect of DOT1L inhibition on cell apoptosis. The percentage of apoptotic cell populations include both early apoptotic cells (Annexin V^+^) and late apoptotic cells (Annexin V^+^/DAPI^+^). *P*-values were calculated using paired t-test. All cell lines were treated for six days with the following concentrations of DOT1Li: 100nM for Oci-Ly7 and SUDHL10; 250nM for SUDHL6; and 2µM for WSU-NHL. Error bars represent the standard error of the mean (SEM) of three biological replicates. ***P-value < 0.001 and *P-value < 0.05. “ns” indicates not significant.

To assess the effect on cellular proliferation, we treated GCB-DLBCL cell lines for six days with different concentrations of DOT1Li (0.1µM – 2µM) to identify the lowest concentration required to maximally reduce H3K79 methylation **(Figure 1B; supplemental Figure 1A, B)**. Accordingly, 100nM DOT1Li was chosen for Oci-Ly7 and SUDHL10; 250nM DOT1Li for SUDHL6 **(Figure 1C)**. The cell line WSU-NHL required 2µM DOT1Li for maximally reducing the level of H3K79me2 after six days. This likely relates to its long doubling time (∼45-57h) **(supplemental Figure 1C)**, since loss of H3K79me2 is mainly determined by passive dilution due to cell proliferation ^25^. H3K79 hypomethylation resulted in a substantial reduction in cell proliferation of DOT1Li-responsive GCB-DLBCL cell lines **(Figure 1D)**. However, a common defect in cell cycle progression and cell survival was not observed **(Figure 1E, F; supplemental Figure 1D, E)**.

### DOT1L and EZH2 are co-dependent and co-inhibition acts synergistically in GCB-DLBCL cell lines

Since the loss of either *Dot1L* or *Ezh2* in mouse B cells prohibits differentiation towards GC B cells, targeting both epigenetic regulators might synergistically impair the fitness of GCB-DLBCL. To explore this hypothesis, we applied STRING clustering analysis to the top 20 genes that are co-dependent with DOT1L based on similar DepMap scores (CRISPR DepMap Public 23Q2 – Chronos Score) **(Figure 2A)**. The subunits of the PRC2 complex (cluster 1) shared a strong co-dependency with DOT1L in over 1000 cell lines. Of the identified PRC2 subunits the co-dependency of EZH2 with DOT1L was more pronounced in DLBCL cell lines as compared to all other cell lines in the DepMap database. This strong co-dependency led us to explore co-inhibition of DOT1L and EZH2 in targeting GCB-DLBCL.

**Figure 2.**
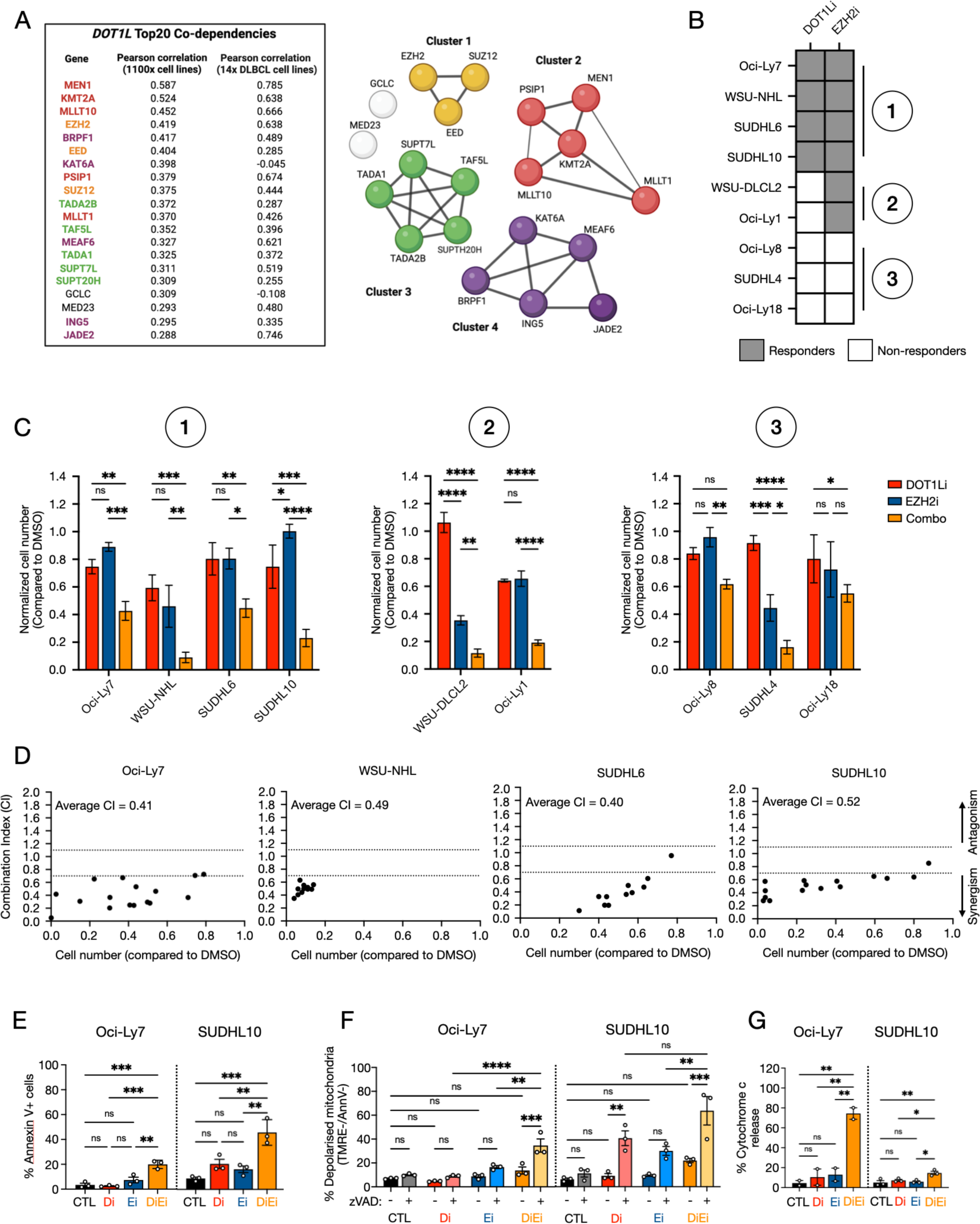
DOT1L and EZH2 are co-dependent and their co-inhibition synergizes in controlling the proliferation of GCB-DLBCL cell lines. (A) Top20 DOT1L co-dependent genes based on Pearson correlation in 1100x cell lines and 14x DLBCL cell lines derived from the DepMap database. STRING analysis revealed four clusters (1) Polycomb Repressor Complex 2 (PRC2), (2) MLL protein complex, (3) SAGA-type co-activator complex, and (4) MOZ/MORF-complex. (B) Schematic overview of GCB-DLBCL cell lines and their responsiveness towards DOT1Li and EZH2i single treatments. (C) Bar graphs represent the effect of DOT1Li alone, EZH2i alone, or the combination of DOT1Li and EZH2i (combo) in a panel of GCB-DLBCL cell lines. Group 1 represents cell lines that were sensitive to both inhibitors, and were treated with EC50-dose of DOT1Li, EC20-dose of EZH2i, or the combination. Group 2 represents cell lines that were only sensitive to EZH2i, and were treated with 250nM DOT1Li and 253 nM EZH2i for WSU-DLCL2; 275 nM DOT1Li and 400 nM EZH2i for Oci-Ly1. Group 3 represents cell lines that were non-responsive to both DOT1Li and EZH2i, and were treated with 100 nM DOT1Li and 5 µM EZH2i for Oci-Ly8; 250 nM DOT1Li and 5 µM EZH2i for SUDHL4; 540 nM DOT1Li and 4.18 µM EZH2i for Oci-Ly18. Error bars represent the standard error of the mean (SEM) of three biological replicates. Cell numbers were measured by flow cytometry after 12 days of treatment. (D) The synergistic potential between DOT1Li and EZH2i for GCB-DLBCL cell lines from group 1. Each datapoint represents a combination of DOT1Li and EZH2, comparing the relative cell number to the Combination Index (CI) score. The CI was calculated using Chou-Talalay scoring methodology in CompuSyn, representing three biological replicates. (E) The combined inhibition of DOT1L and EZH2 induces apoptosis in Oci-Ly7 and SUDHL10 cell lines. Cells were stained with Annexin V/DAPI and measured by flow cytometry. Annexin V^+^ population includes both DAPI^+^ and DAPI^-^ cells. P-values were calculated using one-way ANOVA. (F) Bar graph depicts the percentage of viable cells with depolarized mitochondria (TMRE^-^/AnnV^-^) in Oci-Ly7 and SUDHL10. Apoptosis was inhibited with 25µM pan-caspase inhibitor zVAD from day 3 till day 6. (G) Induction of cytochrome c release by the combination treatment in viable cells. P-values were calculated using one-way ANOVA. For annexin V, TMRE, and cytochrome c measurements, cells were treated for six days with the following conditions, (CTL) <0.01% (v/v) DMSO for Oci-Ly7 and SUDHL10; (Di) 100nM and 60nM of DOT1Li for Oci-Ly7 and SUDHL10, respectively; (Ei) 1000nM and 85nM of EZH2i for Oci-Ly7 and SUDHL10, respectively; and combination of these concentrations for DOT1Li and EZH2i combination treatment (DiEi). For Oci-Ly7, the concentrations of DOT1Li and EZH2i represent the dose wherein H3K79me2 and H3K27me3 are maximally reduced after six days. For SUDHL10, the concentrations are based on the cross-drug design data from day 8, wherein synergy was observed with sufficient cells to be measured. All experiments are performed in three biological replicates, except for cytochrome c release assay of Oci-Ly7 (n=2). Error bars represent the standard error of the mean (SEM). P-values were calculated using two-way ANOVA. ****P-value < 0.0001, ***P-value < 0.001, **P-value < 0.01, *P-value < 0.05. “ns” indicates not significant

To investigate the synergistic potential between DOT1Li and the EZH2 inhibitor (EZH2i) GSK343, we utilized a cross-drug combination design ^26,27^. Dose-response curves provided the EC concentrations of DOT1Li and EZH2i **(supplemental Figure 2A; supplemental Table 2)**. Based on the trend of the dose-response curves, GCB-DLBCL cell lines were divided into three groups of cell lines: (1) sensitive to both inhibitors, (2) non-responsive to DOT1Li but responsive to EZH2i, and (3) non-responsive to both DOT1Li and EZH2i **(Figure 2B)**. In group 1, the combination treatment resulted in a greater reduction in cell number compared to DOT1Li (EC50-dose) and EZH2i (EC20-dose) alone **(Figure 2C)**. For DOT1Li and EZH2i non responsive cell lines without EC values, we used concentrations at which H3K79me2 and H3K27me3 levels were maximally reduced after six days of treatment **(supplemental Figure 2B)**. In group 2, WSU-DLCL2 and Oci-Ly1, showed that adding DOT1Li to EZH2i resulted in a greater loss of cells compared to the single treatment of each inhibitor **(Figure 2C)**. For group 3, the combination treatment sensitized the DOT1Li/EZH2i non-responder SUDHL4, while the effect size was minimal for Oci-Ly8 and Oci-Ly18. For cell lines responsive to both DOT1Li and EZH2i, we applied the Chou-Talalay synergy metric to calculate the combination index (CI) score of multiple combinations of DOT1Li and EZH2i ^28^. Based on the overall CI scores, the combination of DOT1Li and EZH2i acted synergistically (CI < 0.7) in GCB-DLBCL cell lines responsive to both DOT1Li and EZH2i alone **(Figure 2D)**.

To study whether the synergy between DOT1Li and EZH2i induces apoptosis in GCB-DLBCL, we applied flow cytometry to determine the frequency of apoptotic cells. The proportion of Annexin V-positive cells increased in both Oci-Ly7 (from 2.5% to 20%) and SUDHL10 (16.4% to 33%) compared to DMSO **(Figure 2E)**. To determine whether apoptosis is induced by mitochondrial depolarization in GCB-DLBCL cell lines upon DOT1Li, EZH2i, or combination treatment, we inhibited apoptosis using the pan-caspase inhibitor zVAD-fmk (zVAD) and measured the percentage of viable cells with depolarized mitochondria (TMRE^-^, Annexin V^-^) **(Figure 2F; supplemental Figure 2C)**. Upon the combination treatment, zVAD decreased the early apoptotic population (AnnV^+^), which resulted in a significant increase in a population with depolarized mitochondria, from 10% to 29% in Oci-Ly7, and 25% to 70% in SUDHL10. The combination treatment was associated with mitochondrial depolarization, increased cytochrome c release, and caspase-mediated apoptosis **(Figure 2G)**. These insights imply that mitochondrial-depolarization is a main driver of apoptosis upon combination treatment in the tested GCB-DLBCL cell lines.

Taken together, these data validate a strong dependency of GCB-DLBCLs on the activity of DOT1L and EZH2, and co-inhibition impaired the fitness in seven out of nine GCB-DLBCL cell lines tested.

### DOT1L and EZH2 cooperatively repress PRC2 targets to maintain the GC B cell identity in DLBCL

Previous work by Aslam et al ^17^ suggested that DOT1L indirectly represses PRC2 target genes in murine B cells by activation of *Ezh2*. To address whether this functional relationship persists in human GCB-DLBCL and relates to combination treatment sensitivity, gene expression and protein levels of EZH2 were determined in two sensitive cell lines, Oci-Ly7 and SUDHL6, and two non-responsive cell lines, Oci-Ly8 and Oci-Ly18. For each cell line, we chose concentrations of DOT1Li and EZH2i wherein H3K79me2 and H3K27me3 were maximally reduced, respectively, to compare the response between cell lines and reduce potential off-target effects. Due to the bimodal distribution of H3K27me3 peaks in SUDHL6 upon EZH2i, a concentration of 2 µM EZH2i was chosen which resulted in substantial decrease in proliferation in combination with DOT1Li when compared to EZH2i alone **(supplemental Figure 3A)**. Concentrations of DOT1Li required for the loss of H3K79me2 neither transcriptionally reduced EZH2 mRNA levels nor affected EZH2 protein expression **(Figure 3A; supplemental Figure 3B)**. Globally, H3K27me3 levels were not affected upon DOT1Li, indicating that DOT1L does not affect the activity of EZH2 or the PRC2 complex **(Figure 3B)**.

**Figure 3.**
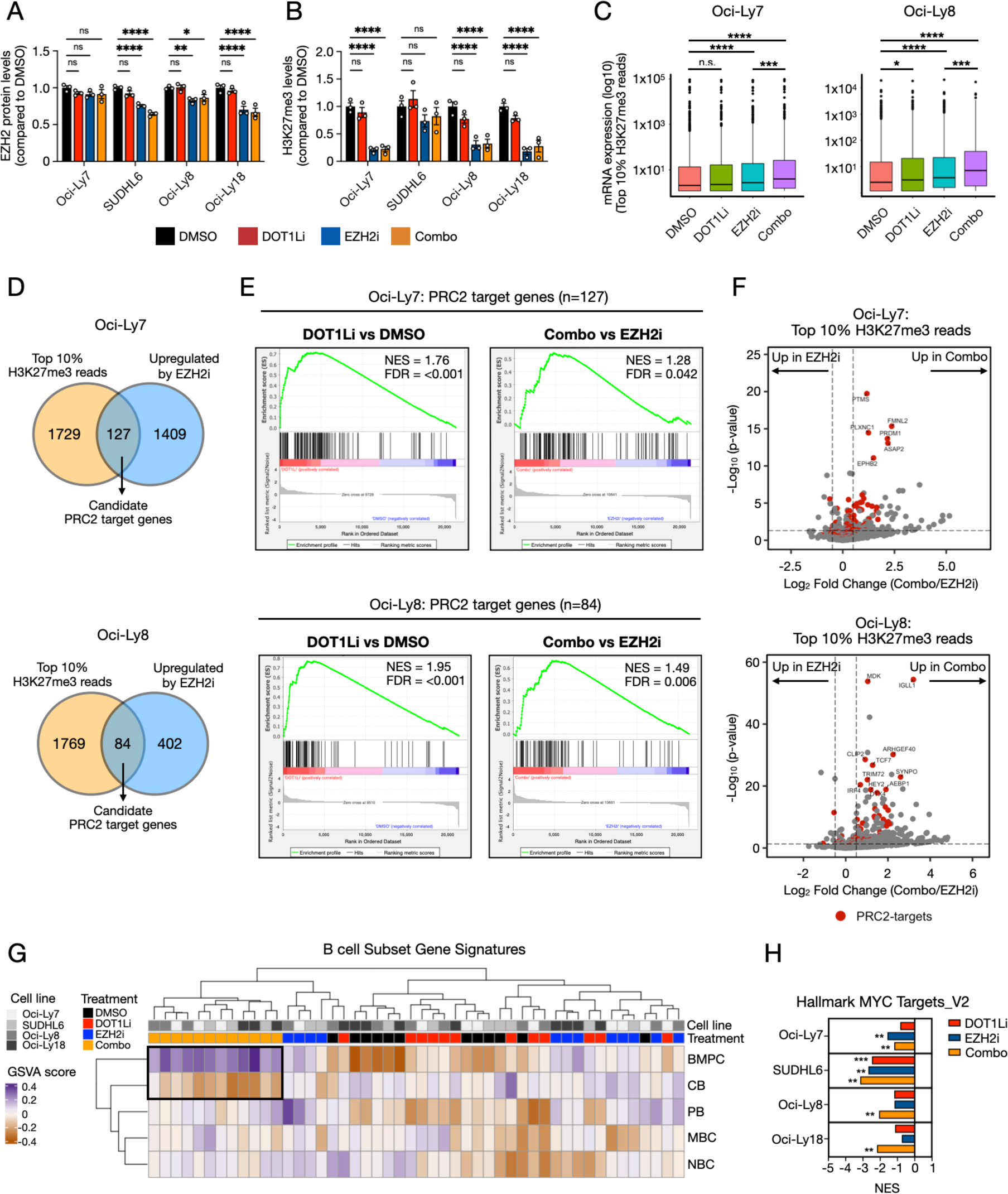
Inhibition of DOT1L and EZH2 promotes GCB-DLBCL differentiation into plasma-like cells. (A) EZH2 protein levels as measured by flow cytometry in four independent GCB-DLBCL cell lines treated with DMSO, DOT1Li alone, EZH2i alone, or combination of both treatments. (B) H3K27me3 levels measured by flow cytometry. (C) Box plot showing the expression changes of the top 10% protein-coding genes with H3K27me3 reads from transcription start site (TSS) to +3kb. Significance was calculated using Wilcoxon rank sum test. (D) Venn-diagram displays the overlap between the top 10% H3K27me3 read counts from TSS to +3kb of protein-coding genes and significantly upregulated genes upon EZH2 inhibition in Oci-Ly7 and Oci-Ly8 cell lines. Overlapped genes are defined as candidate PRC2 target genes. Oci-Ly7: average H3K27me3 read counts = 0.116; Oci-Ly8: average H3K27me3 read counts = 0.156. (E) Significant enrichment of PRC2 targets in DOT1Li-treated cell lines versus DMSO and combo-treated cell lines versus EZH2i alone based on GSEA. (F) Volcano plot represents the differential expression of the top 10% of protein-coding genes with H3K27me3 reads between combo treatment versus EZH2i alone. The highlighted genes (red) represent the defined candidate PRC2 target genes within the top 10% for both Oci-Ly7 and Oci-Ly8. (G) Unsupervised clustering of GSVA scores of five B cell subset gene signatures based on RNA sequencing data from four independent GCB-DLBCL cell lines, treated with DOT1Li, EZH2i, or the combination of both treatments. Cell lines treated with combination clustered together based on the gain of a bone marrow plasma cell (BMPC) signature and the loss of their centroblast (CB) signature. (H) GSEA identified the significant depletion of MYC target genes by the combination treatment compared to DMSO. Cells were treated for six days with the following concentrations of DOT1L1: 100nM for Oci-Ly7 and Oci-Ly8; 250 nM for SUDHL6; and 2µM for Oci-Ly18, and the following concentrations of EZH2i: 1µM for Oci-Ly7; 5µM for Oci-Ly8 and Oci-Ly18; and 2µM for SUDHL6. DOT1Li and EZH2i concentrations are based on the doses wherein H3K79me2 and H3K27me3 are maximally reduced. NBC = naïve B cells; CB = centroblasts; PB = plasmablasts; MBC = memory B cells; BMPC = bone marrow plasma cells. Error bars represent the standard error of the mean (SEM) of three biological replicates. P-values were calculated using two-way ANOVA. ****P-value < 0.0001, ***P-value < 0.001, **P-value < 0.01, *P-value < 0.05. “ns” indicates not significant. The false discovery rate (FDR) was calculated using GSEA. ***q-value < 0.001, ** q-value <0.01.

To determine whether DOT1L promotes repression of PRC2 target genes independently of EZH2, we performed H3K27me3 ChIP-seq on untreated responder Oci-Ly7 and non-responder Oci-Ly8 cell lines, and selected the top 10% of protein-coding genes with the highest H3K27me3 read counts from transcription start site + 3 kilobases for subsequent analysis. Interestingly, both GCB-DLBCL cell lines showed significant derepression of H3K27me3-containing genes upon combination treatment when compared to EZH2i alone **(Figure 3C)**. To identify the candidate direct targets of PRC2, we overlapped all the genes that were significantly upregulated upon EZH2i with the genes that contain the top 10% H3K27me3 read counts for both cell lines **(Figure 3D)**. We used this list of PRC2-target genes for gene set enrichment analysis (GSEA) and observed that DOT1Li was able to significantly derepress candidate PRC2 target genes in both Oci-Ly7 and Oci-Ly8 when compared to DMSO **(Figure 3E; supplemental Figure 3C, D)**. Moreover, this effect of DOT1Li was also observed upon combination treatment, indicating that the effect of DOT1L in repressing PRC2 targets might indeed be independent of EZH2. Along with the fact that DOT1L-mediated H3K79 methylation is generally associated with gene activation, suggests that DOT1L indirectly regulates the repression of PRC2 target genes.

In the absence of *Dot1l* or *Ezh2,* GC B cells cannot maintain their identity and instead differentiate into a dysfunctional plasma cell-like state ^11,17,18,29^. Based on cooperativity of DOT1L and EZH2 in repressing PRC2 target genes in GCB-DLBCL, we investigated whether the combined inhibition of DOT1L and EZH2 enhances plasma cell differentiation. Interestingly, among the derepressed PRC2 targets we identified essential genes for B cell differentiation, including *PRDM1* in Oci-Ly7, and *IRF4* in Oci-Ly8, as well as genes associated with cellular morphogenesis, including *ASAP2* and *FMNL2* in Oci-Ly7, and *SYNPO* and *CLIP2* in Oci-Ly8 **(Figure 3F)**. As an independent approach, we first established differentiation-state specific signatures unique for NBC, CB, MBC, PB, and BMPC using the GenomicScape database. Unsupervised clustering of the GSVA scores revealed that all the cell lines treated with the combination treatment were clustered together and gained a BMPC signature at the expense of a CB-signature **(Figure 3G)**. This cooperative activity of the epigenetic writers DOT1L and EZH2 with opposing transcriptional effects in maintaining the identity of GCB-DLBCL was confirmed independently using published gene signatures **(supplemental Figure 3E)** ^11,30^. This crosstalk between DOT1L and EZH2 appeared to be conserved among different GCB-DLBCL cell lines and was independent of their responsiveness to the treatments.

Exploring the CB and BMPC signatures, these gene sets/signatures were enriched for the pathways that are associated with the pro-proliferative capacity of GC B cells and the massive morphogenic and migratory alterations associated with PC differentiation, respectively **(supplemental Figure 3F)**. Within CB signatures gene sets, we found that upon the combination treatment, replication-associated pathways were significantly downregulated, correlating with their reduced proliferative capacity **(supplemental Figure 3G)**. PC differentiation is associated with the upregulation of MAPK and IL4/IL13 signaling pathways, which are known to directly contribute to the production of immunoglobulins by PCs and induce *PRDM1* (BLIMP-1) expression, a key PC transcription factor ^31,32^. Indeed, the tendency of GCB-DLBCL to adopt a PC-like identity was associated with the upregulation of *PRDM1* and downregulation of B cell transcription factors *FOXO1* and *IRF8* **(supplemental Figure 3H)**.

Besides the dependency on DOT1L and EZH2 activity, the transcriptional regulator MYC is an essential initiator and driver of the GC-reaction and -identity ^33^. The fact that the GCB-DLBCLs lose their GC identity in the absence of DOT1L and EZH2 activity led us to determine whether this also coincides with specific alterations in MYC target gene expression. Interestingly, and in line with the loss of GC identity and gain of PC identity upon the combination treatment, analysis of *MYC*-translocated GCB-DLBCL cell lines revealed that MYC target genes were significantly downregulated upon combination treatment **(Figure 3H)**. These findings indicate a strong dependency on the epigenetic writers DOT1L and EZH2 in maintaining MYC target gene activity in *MYC* rearranged GCB-DLBCLs.

### Combined inhibition of DOT1L and EZH2 prevents outgrowth of a human DLBCL xenograft

To determine the potency of a combined epi-drug based treatment in controlling human DLBCL *in vivo*, we xeno-grafted the Oci-Ly7 cell line in immunodeficient NSG mice. Mice were randomized to vehicle, DOT1Li, EZH2i, or the combination treatment (Combo). Based on its favorable pharmacokinetic properties, we chose compound 11 (DOT1L-IN-5) as DOT1L inhibitor ^34,35^. As FDA-approved inhibitor, we opted for tazemetostat as the EZH2 inhibitor. In comparison to vehicle, DOT1L-IN-5 or tazemetostat alone, the Combo treatment prevented the tumor outgrowth in all the mice and completely eradicated the tumor in four out of seven mice after 24 days of treatment, as measured by calipers (Combo vs DOT1L-IN-5, *P* < 0.001; Combo vs tazemetostat, *P* < 0.001) **(Figure 4A)**. No significant difference in restricting tumor outgrowth was observed between DOT1L-IN-5 and tazemetostat treatments (DOT1L-IN-5 vs tazemetostat, *P* = 0.8172). The simultaneous reduction in the levels of H3K79me2 and H3K27me3 in PBMCs upon Combo treatment indicates the efficacy of both drugs *in vivo,* which was necessary for the observed tumor growth control **(Figure 4B, C)**. Histological sections of the xenografts revealed that the Combo treatment promoted apoptosis in the tumor cells and necrosis around the tumor area **(supplemental Figure 4A, B)**. By transforming the non-linear model of tumor volume to log-scaled linear model, we identified that the Combo treatment had a significant synergistic effect on tumor control when compared to the single treatments (*P* = 0.0023) **(Figure 4D)**. While the median tumor survival for mice treated with vehicle, DOT1L-IN-5, and tazemetostat was 12, 17, and 19 days, respectively, all mice treated with Combo reached the fixed endpoint of 24 days, indicating the synergy of the two epi-drugs in preventing tumor outgrowth and prolonging survival **(Figure 4E)**.

**Figure 4.**
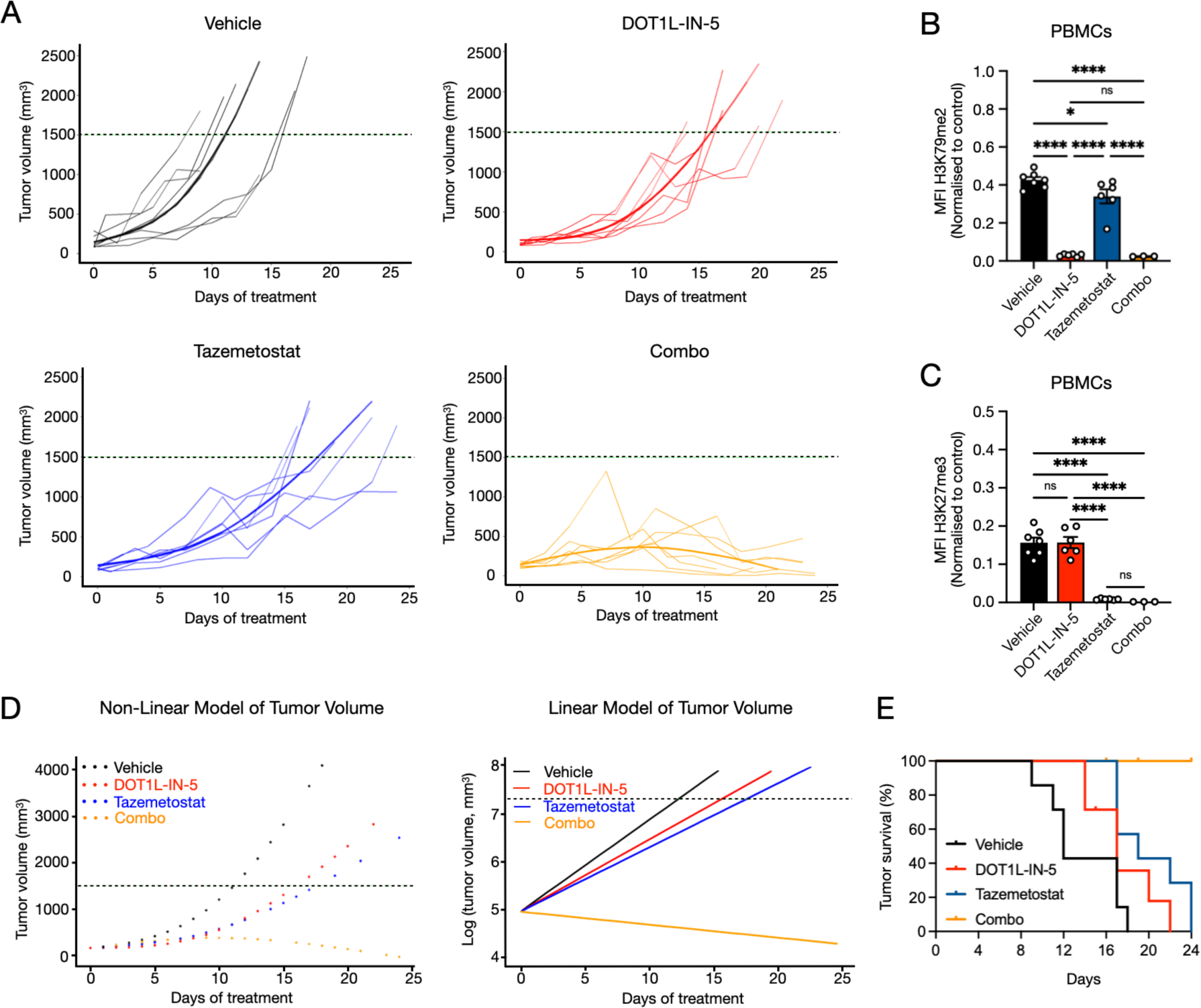
Human GCB-DLBCL xenograft: Combined inhibition of DOT1L and EZH2 prevents tumor outgrowth. (A) Immunodeficient NSG mice xenografted with Oci-Ly7 were treated twice daily with vehicle, 75mg/kg DOT1L-IN-5, 300mg/kg tazemetostat, or a combination (n=7 mice per treatment group). A natural cubic spline of the second order was used to model the mean tumor growth over time, depicted by the bold line. (B-C) Median fluorescence intensity (MFI) of H3K79me2 and H3K27me3 levels cells from *ex vivo* peripheral blood mononuclear cells (PBMCs) after terminal bleeding. MFI is normalized to *in vitro* cultured Oci-Ly7 as control. Error bars represent the standard error of the mean (SEM). P-values were calculated using one-way ANOVA. (D) Linear mixed model on log transformed scale of tumor volume has a significant synergistic effect by the combination treatment (*P* = 0.0023). (E) Kaplan-Meier curve comparing tumor survival in percentage of vehicle, DOT1L-IN-5, tazemetostat, and the combination in treated mice. DOT1L-IN-5 vs tazemetostat, *P* = 0.2074; DOT1L-IN-5 vs Combo, *P* < 0.001; tazemetostat vs Combo, *P* = 0.0042; Combo vs Vehicle, *P* < 0.001 ****P-value < 0.0001 and *P-value < 0.05. “ns” indicates not significant.

To our knowledge, the toxicity of this combination treatment has not yet been described. The overall body weight remained stable, with one exception **(supplemental Figure 4C)**. Histopathological examination of the bone marrow and spleen revealed a great reduction in erythropoiesis based on the depletion of erythropoietic cells, a side effect not observed in the other cohorts **(supplemental Figure 4D)**. Unexpectedly, three out of seven mice treated with tazemetostat alone acquired DLBCL localization in the bone marrow, which was not observed in other treatment cohorts **(supplemental Figure 4E)**. Histopathology of the kidneys provided indications of nephrotoxicity such as renal degeneration, necrosis, and hemorrhages in a few mice especially within the DOT1L-IN-5 (4 out of 7 mice) and Combo-treated (2 out of 5 mice) cohorts **(supplemental Figure 4F)**. No toxicity was observed in the liver, gastrointestinal tract, heart, lungs, and brain.

## Discussion

Approximately 40% of DLBCL cases are refractory to or relapse by standard-of-care R-CHOP therapy, with limited alternative therapeutic options available ^36,37^. GCB-like DLBCL arises from GC B cells and closely resembles their cell of origin. Given the strong dependency of GC B cells on DOT1L and EZH2 in establishing and maintaining their pro-proliferative identity, we explored the role of these epigenetic writers as potential therapeutic targets for GCB-DLBCLs. Our data supports a well-preserved role for DOT1L and EZH2 in cooperatively establishing an epigenetic barrier necessary for the maintenance of the GC B cell identity in MYC-rearranged DLBCL. Combined targeted inhibition promotes the transition from a GC B cell phenotype into a ‘so-called’ plasma cell-like state with compromised cell survival. Most (7 out of 9) of the GCB-DLBCL cell lines tested *in vitro* were responsive towards the combined inhibition of DOT1L and EZH2, confirming the co-dependency of GCB-DLBCL on these epigenetic writers. Importantly, the same synergistic effect was observed *in vivo* for GCB-DLBCL cell line xenografts of Oci-Ly7.

In this proof-of-concept study, we evaluated the effectiveness and toxicity of the combination therapy *in vivo.* In support of our *in vitro* results, we demonstrate that targeting of both DOT1L and EZH2 effectively controls tumor growth. Pathologically, the combination treatment primarily suppressed erythropoiesis in the bone marrow and spleen, potentially leading to anemia, a side effect warranting further investigation ^38^. Anemia is a common and mild side effect of R-CHOP therapy in DLBCL patients, typically manageable with blood transfusions ^38,39^. Future *in vivo* studies are needed to identify the optimal inhibitors, aiming to minimize therapy-related erythropoiesis loss and explore its link to erythropoietin (Epo) production by the kidneys ^40^. Regardless of these side effects, our findings unveil combined epi-drugging as a promising therapeutic opportunity for targeting GCB-DLBCL.

It is well established that PRC2, in cooperation with Polycomb Repressor Complex 1 (PRC1), is essential for maintaining cellular identity and regulating cell-fate decisions by repressing its target genes. Dysregulation of PRC2, leading to altered expression of target genes, can promote phenotypic plasticity in multiple cell types ^41,42^. Studies have shown that EZH2, the catalytic subunit of PRC2, epigenetically silences genes associated with cell cycle checkpoints and plasma cell differentiation in both GC B cells and GCB-DLBCL, highlighting its fundamental and translational value ^29^. Although the loss of activity of either DOT1L or EZH2 suffices to derepress PRC2 target genes to some extent, neither can individually shape the molecular landscape of the cell required to obtain the plasma cell-like state. Instead, the combined inhibition of DOT1L and EZH2 enhances the derepression of key PRC2 targets, including PC transcription factor *PRDM1*, to such an extent that GCB-DLBCLs undergo the partial transition into PCs.

This finding suggests that, although molecularly DOT1L and H3K79me are associated with transcriptional activation of genes, DOT1L supports the repression of PRC2 target genes independently of EZH2 in GCB-DLBCL. This observation hints at the existence of an alternative repressive mechanism by which DOT1L indirectly represses PRC2 target genes, considering the major dependency of PRC2 on EZH2 activity. Sparbier et al. reported that even in the absence of H3K27me3, gene silencing in bivalent promoters is still accomplished through the deposition of H2AK119ub1 by PCGF1, a subunit of non-canonical PRC1.1 ^43^. This indicates that both H2AK119ub1 and H3K27me3 control the transcription of PRC2 target genes, and their complete derepression requires the loss of these epigenetic modifications. However, it remains to be elucidated whether DOT1L might be crucial either in directly or indirectly regulating PRC1 and H2AK119ub1 deposition at PRC2 target genes.

Corrupted cellular differentiation is a hallmark in the development of many types of malignant tumors ^44^. Here, we propose a differentiation-based therapeutic strategy targeting DOT1L and EZH2 to epigenetically drive the differentiation of GCB-DLBCL into a terminally differentiated plasma cell-like state. This differentiated state is associated with the loss of MYC target gene expression and decreased cellular survival. Our findings not only highlight the role of DOT1L and EZH2 in maintaining the MYC-driven GC B cell identity in DLBCL through the cooperative repression of PRC2 target genes but also introduce a novel therapeutic approach for DLBCL treatment.

## Acknowledgment

We thank the flow cytometry core facility, mouse preclinical intervention facility, genomic core facility, and experimental animal pathology facility at the Netherlands Cancer Institute for advice and support. We thank T. van den Brand for helpful discussions in ChIP-seq analysis. We highly appreciate Dr. J.E.J. Guikema for providing the DLBCL cell lines and for critically reading of the manuscript. This research was supported by an institutional grant of the Dutch Cancer Society and of the Dutch Ministry of Health, Welfare and Sports. This work was supported and made possible by the Dutch Cancer Society KWF grant no. 12825 to H.J. and F.v.L. The funders had no role in study design, data collection and interpretation, or the decision to submit the work for publication.

## Authorship

Contributions: H.J. and F.v.L. initiated and designed the study. C.G. and R.N. conducted most of the experiments and data analysis. B.M. established and optimized the semi-automated robotics protocol. J.T. performed pre-processing of ChIP-seq experiments and provided expertise in both ChIP-seq and RNA-seq. J.S. performed the animal pathology analysis. L.A. performed all *in vivo* tumor control statistics. D.d.G. provided expertise in RNA-seq analyses. M.H.P.d.G., J.J., E.K., and M.K. performed experiments and analyzed the data. M.A.A. initiated the project, helped with setting up the initial experiments, RNA seq analysis and provided critical expertise. C.G., R.N., F.v.L., and H.J. wrote the manuscript.

## Supplemental materials

### Cell lines

GCB-DLBCL cell lines were obtained as follows: SUDHL4 and WSU-DLCL2 were purchased from DSMZ, German Collection of Microorganisms and Cell cultures GmbH. The others cell lines, Oci-Ly1, Oci-Ly7, Ocy-Ly8, Oci-Ly18, SUDHL6, SUDHL10, WSU-NHL, were kindly provided by Dr. Jeroen Guikema (The Academic Medical Center, AMC) and Blanca Scheijen (The Radboud University Medical Center, Nijmegen). All GCB-DLBCL cell lines were cultured in Iscove’s Modified Dulbecco Medium (IMDM) (Thermofisher scientific) supplemented with 8% fetal cow serum (Capricon Scientific), 1% GlutaMAX (Gibco), and penicillin/streptomycin (Sigma-Aldrich). Cells were maintained at 37°C and 5% CO_2_.

### Cell cycle analysis

Cells were washed and resuspended in PBS (2×10^5 cells/100ul). Cells were fixed with ethanol to a final concentration of 70%, and stored at 4°C for at least 2 hours. Cells were then resuspended in 200ul of PBS and after 60 seconds, centrifuged for 3 minutes at 2000 rpm. The cell pellet was resuspended in 200ul of 1:100 DAPI (stock concentration of 70µg/mL). After a 30-minute incubation in the dark at room temperature, samples were acquired by flow cytometry.

### Cytochrome C release

To determine the concentration of digitonin required for permeabilizing the plasma membrane without affecting mitochondrial membrane integrity, we titrated the digitonin concentration (0–100 µg/ml) and incubated the cells with 1:125 TMRE (10 µM) for 30 minutes at 37°C to determine the concentration at which the cells still maintain an active mitochondrial membrane potential.

For the cytochrome c release assay, we stained the cells with LiveDead-Yellow (Invitrogen™, cat. No. L34959) at a dilution of 1:1000 in cold PBS for 15 minutes at 4°C, following the manufacturer’s protocol. The cells were then permeabilized with digitonin (10 µg/ml in Oci-Ly7 and 6 µg/ml in SUDHL10) for 10 minutes at 4°C and fixed with 3.7% formaldehyde in PBS for 20 minutes at room temperature. Subsequently, the cells were stained with Alexa Fluor® 488 anti-Cytochrome c antibody (Biolegend, cat. No. 612308) at a dilution of 1:200 in perm/wash buffer (BD Pharmingen™ Transcription Factor Buffer Set) for 30 minutes at 4°C, followed by two washes with perm/wash buffer. Between washes, the cells were incubated for 10 minutes at 4°C to allow unbound cytochrome c to diffuse passively out of the mitochondria. The cytochrome c negative population represents the cells that have released cytochrome c into the cytosol.

### Gene signature analysis

Pathway enrichment analysis of B cell subset gene sets was performed using g:profiler (https://biit.cs.ut.ee/gprofiler/gost). Gene set enrichment analysis (GSEA) was performed using the GSEA software developed by the Broad Institute (https://www.gsea-msigdb.org/gsea/index.jsp) ^57^.

Enrichment scores generated by GSVA scores the enrichment of each B cell subset gene signature by comparing the expression levels of genes within the gene set to the expression levels of all other genes within each sample. A positive GSVA score suggests that the genes in the gene set are collectively upregulated, indicating the enrichment of a B cell subset gene signature. A negative GSVA score suggests that the genes in the gene set are collectively downregulated, indicating potential suppression of a gene signature.

### Animal handling

All procedures were performed in accordance with Dutch law and the institutional committees (Animal experimental committee and Animal welfare body) overseeing animal experiments at the Netherlands Cancer Institute, Amsterdam.

Mice were housed under standard conditions, including feeding, light cycles, and temperature, with ad libitum access to food and water. All mice were kept in disposable cages in the Laboratory Animal Center (LAC) of the Netherlands Cancer Institute, which minimized the risk of cross-infection, improved ergonomics, and obviated the need for a robotics infrastructure for cage-washing.

### Histochemistry

The full tumor dissection, including surrounding normal tissue, involved harvesting the following organs: GI Tract, kidney, liver, hindleg, spleen, lung, brain, and heart. These tissues were then fixed in 10% formalin (3.7% formaldehyde in water with 1% methanol) for 48 hours. Subsequently, the fixed tumor and organs were embedded in paraffin, and sections of 2 µm thickness were prepared. Finally, the sections were stained with hematoxylin and eosin (H&E).

### Statistical analysis of tumor control in vivo

Statistical analyses were conducted using GraphPad Prism v.10 and R version 4.4.2 (R Development Core Team and the R Foundation for Statistical Computing). Open-source packages such as ‘nlme’ and ‘JMbayes2’, along with packages from the ‘tidyverse’ (including ‘dplyr’, ‘tidyr’, and ‘ggplot’), were integrated into the analysis.

Both linear and nonlinear mixed effect models were employed to analyze repeated measurements of tumor volumes from the same animals over time. These models account for induced correlations among repeated measurements. Natural splines were utilized to fit a nonlinear profile of tumor volumes in individual tumors over time. Time was specified as a continuous covariate, its effect treated as a random effect, and mouse ID as grouping factors. Additionally, the interaction effect of treatment with time was considered a fixed effect. Baseline treatment effects were not considered due to randomization.

Furthermore, a joint model was used to compare treatment groups, utilizing both time-to-event data and tumor growth over time. This approach optimally leverages observed data and enhances statistical power to detect effects. Survival functions for time-to-event outcomes were estimated using the Kaplan– Meier method, with P-values reflecting the Log-rank Mantel-Cox test.

---

**Supplementary Table 1.** Main molecular alterations in GCB-DLBCL cell lines.

| Cell line | Oci-Ly7 | WSU-NHL | SUDHL6 | SUDHL10 | WSU-DLCL2 | Oci-Ly1 | Oci-Ly8 | SUDHL4 | Oci-Ly18 | References |
| --- | --- | --- | --- | --- | --- | --- | --- | --- | --- | --- |
| COO subtype | GCB | GCB | GCB | GCB | GCB | GCB | GCB | GCB | GCB |  |
| Sensitive to DOT1Li | Yes | Yes | Yes | Yes | No | No | No | No | No |  |
| MYC-translocation | + | - | + | + | - | amp | + | + | + | [45-48] |
| BCL2-translocation | - | + | + | + | + | amp | + | + | + | [45, 47] |
| EZH2 | wt | wt | mut | mut | mut | mut | wt | mut | wt | [49, 50] |
| TP53 | mut | mut | mut | mut | mut | mut | mut | mut | mut | [45] |
| CREBBP | wt | mut | mut | mut | n.d. | wt | mut | wt | wt | [51-54] |
| EP300 | wt | wt | mut | mut | wt | n.d. | wt | wt | wt | [52, 54] |
| KMT2D | wt | wt | mut | mut | wt | mut | wt | wt | mut | [55-56] |

**Supplementary Table 2.**
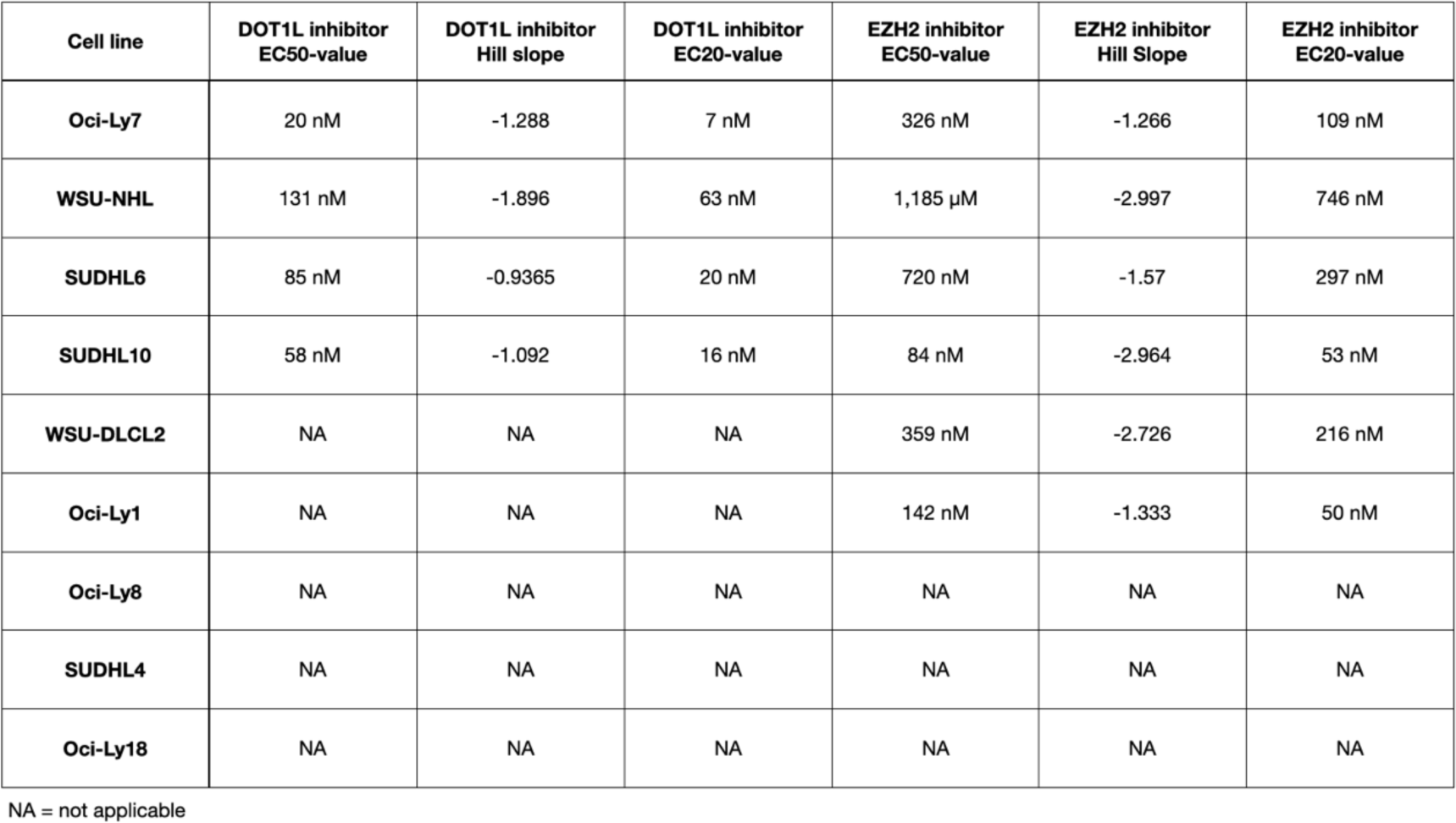
EC-values based on dose response curves of DOT1L and EZH2 inhibitors.

**Supplementary Figure 1.**
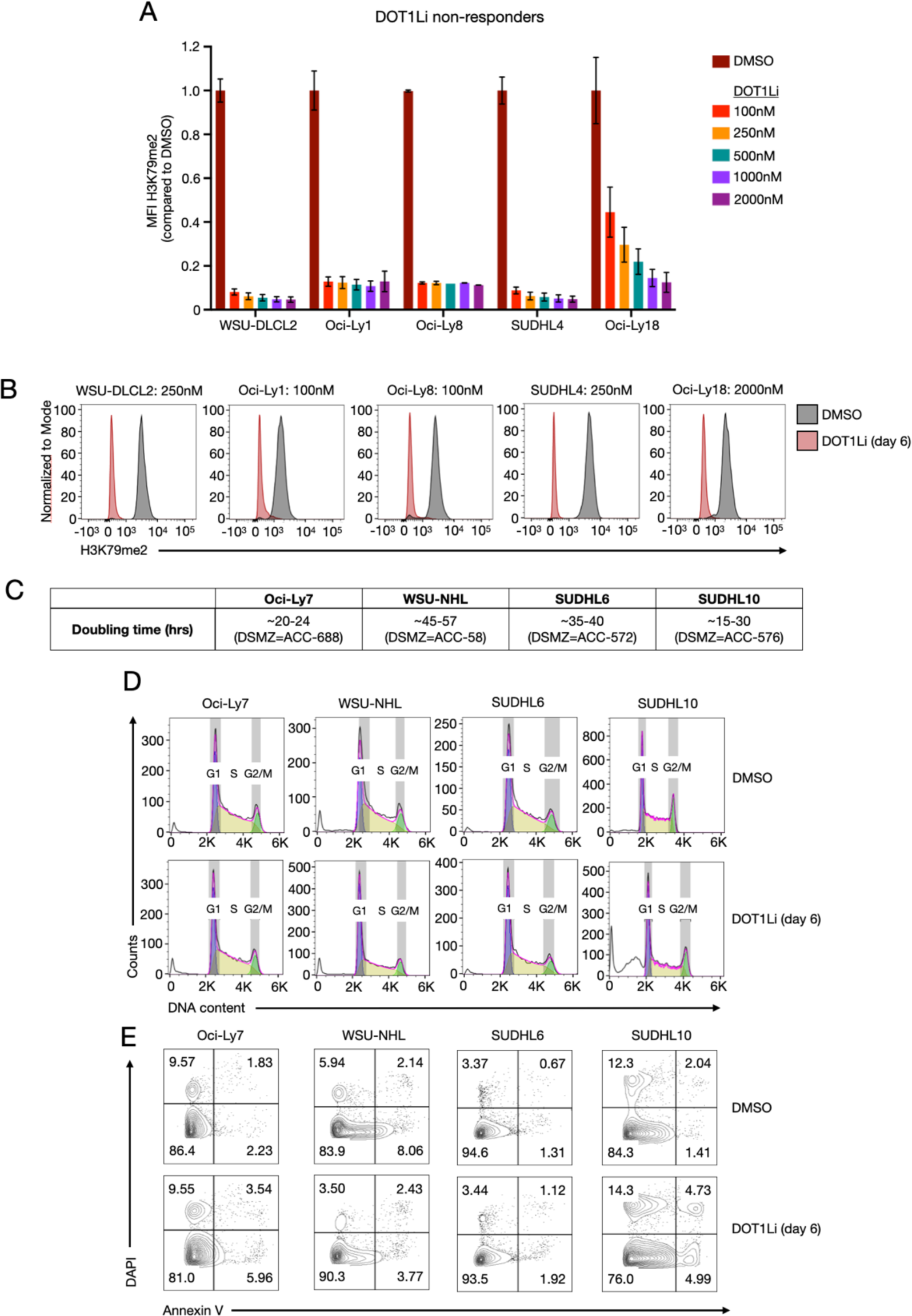
Inhibition of DOT1L affects proliferation and cell viability in DOT1Li-sensitive GCB-DLBCL cell lines. (A) Median fluorescence intensity (MFI) of H3K79me2 upon six days of EPZ-5676 treatment in DOT1Li non-responsive GCB-DLBCL cell lines. Results represent the data from two biological replicates. (B) Representative histograms of the relative intracellular density for H3K79me2 in DOT1Li non-responsive GCB-DLBCL cell lines. Cells were treated for six days with EPZ-5676. (C) The estimated growth rate and doubling time for DOT1Li-sensitive GCB-DLBCL cell lines based on online public data from Leibniz Institute DSMZ. (D) Representative flow cytometry histograms of cell cycle distribution (G1 = purple, S = yellow, and G2 = green) in DOT1Li-sensitive GCB-DLBCL cell lines treated with EPZ-5676. (E) Representative flow cytometry plots of the gating strategy to examine apoptosis of DOT1Li-sensitive GCB-DLBCL cell lines treated with EPZ-5676. Cells were stained with Annexin V/DAPI. GCB-DLBCL cell lines were treated for six days with the following concentrations of EPZ-5676 wherein H3K79me2 was maximally reduced: 100nM for SUDHL10, Oci-Ly7, Oci-Ly1 and Oci-Ly8; 250nM for SUDHL6, WSU-DLCL2, and SUDHL4; 2µM for WSU-NHL and Oci-Ly18.

**Supplemental Figure 2.**
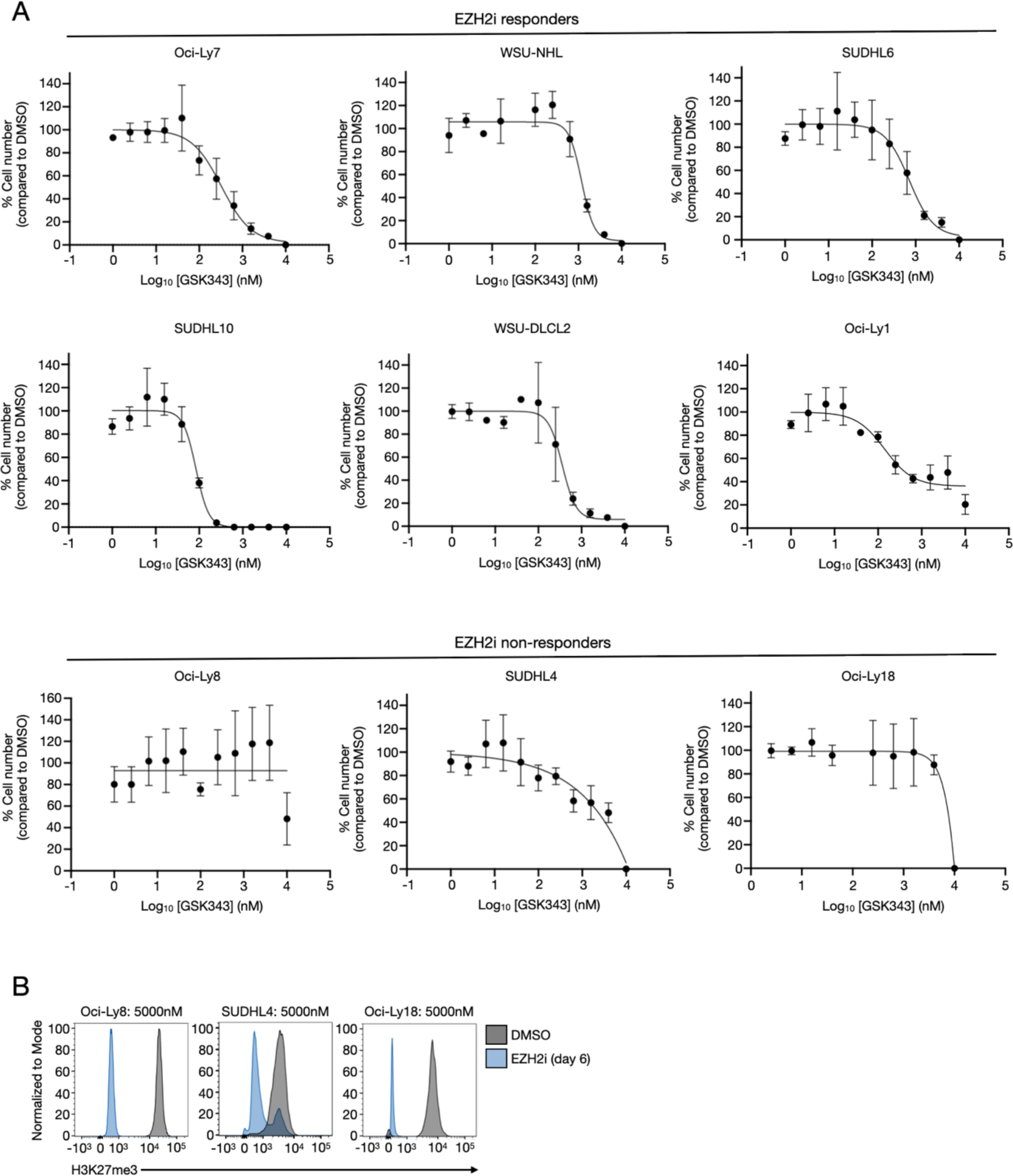
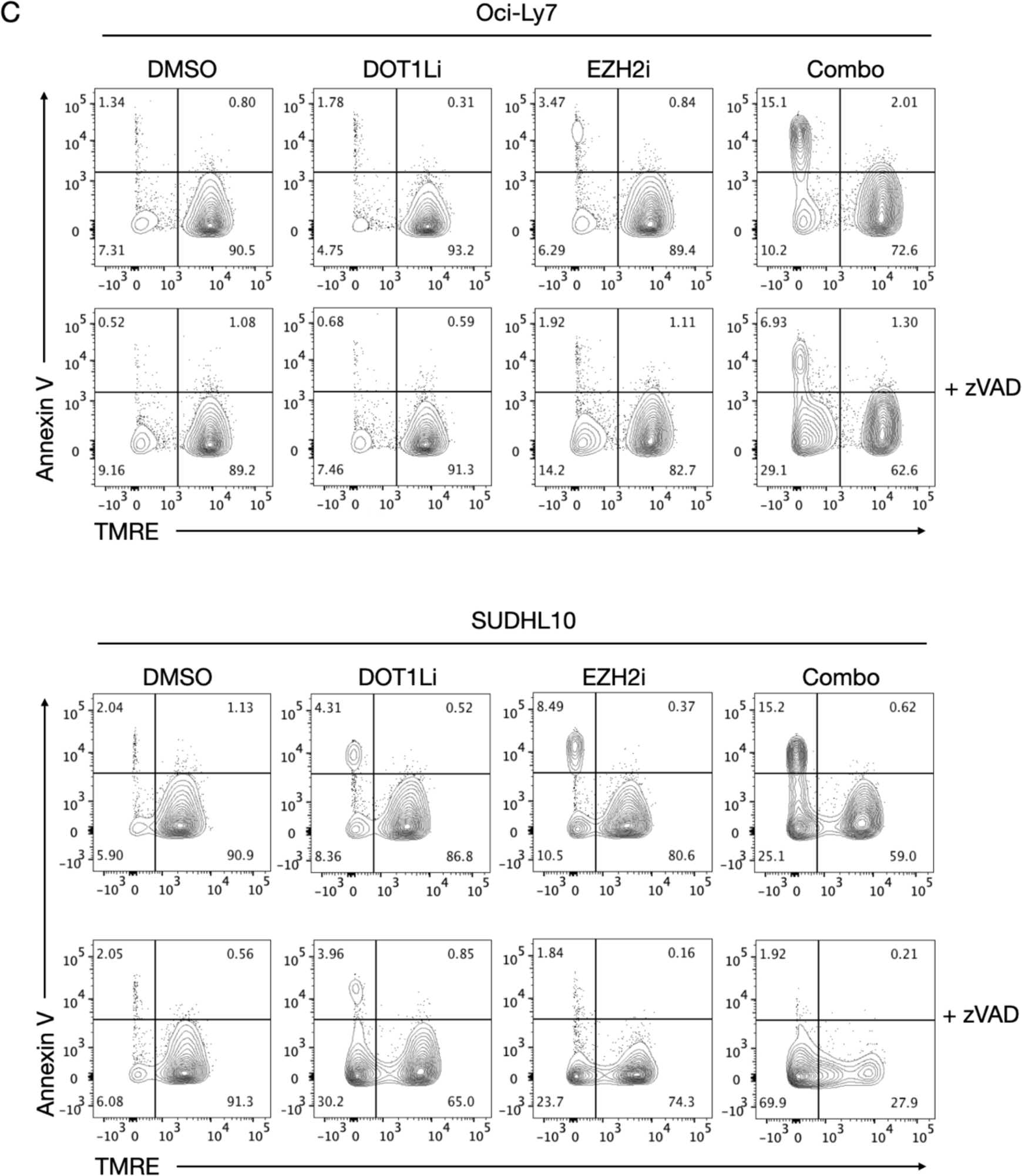
DOT1Li- and EZH2i-responsive GCB-DLBCL cell lines undergo mitochondrial-dependent apoptosis upon combination treatment. (A) Dose-response curves of nine GCB-DLBCL cell lines to EZH2 inhibitor GSK343 after 12 days of treatment. Error bars represent the standard error of the mean (SEM) of three biological replicates. (B) Representative histograms of the relative intracellular density for H3K27me3 in EZH2i non-responsive GCB-DLBCL cell lines treated for six days with 5µM GSK343 where H3K27me3 is maximally reduced. (C) Representative flow cytometry plots showing gating strategy to identify cells population with depolarized mitochondria (TMRE^-^/AnnV^-^) in Oci-Ly7 and SUDHL10. Treatment with the pan-caspase inhibitor zVAD-fmk (zVAD) increased the cell population with depolarized mitochondria upon combination treatment in both cell lines. Oci-Ly7 and SUDHL10 were treated with either DOT1Li and/or EZH2i for six days with the following concentrations: 100nM DOT1Li and/or 1µM EZH2i for Oci-Ly7; 60nM DOT1Li and/or 85nM EZH2i for SUDHL10. Cells were treated with 25µM zVAD or with 0.25% (v/v) DMSO from day 3 to day 6.

**Supplemental Figure 3.**
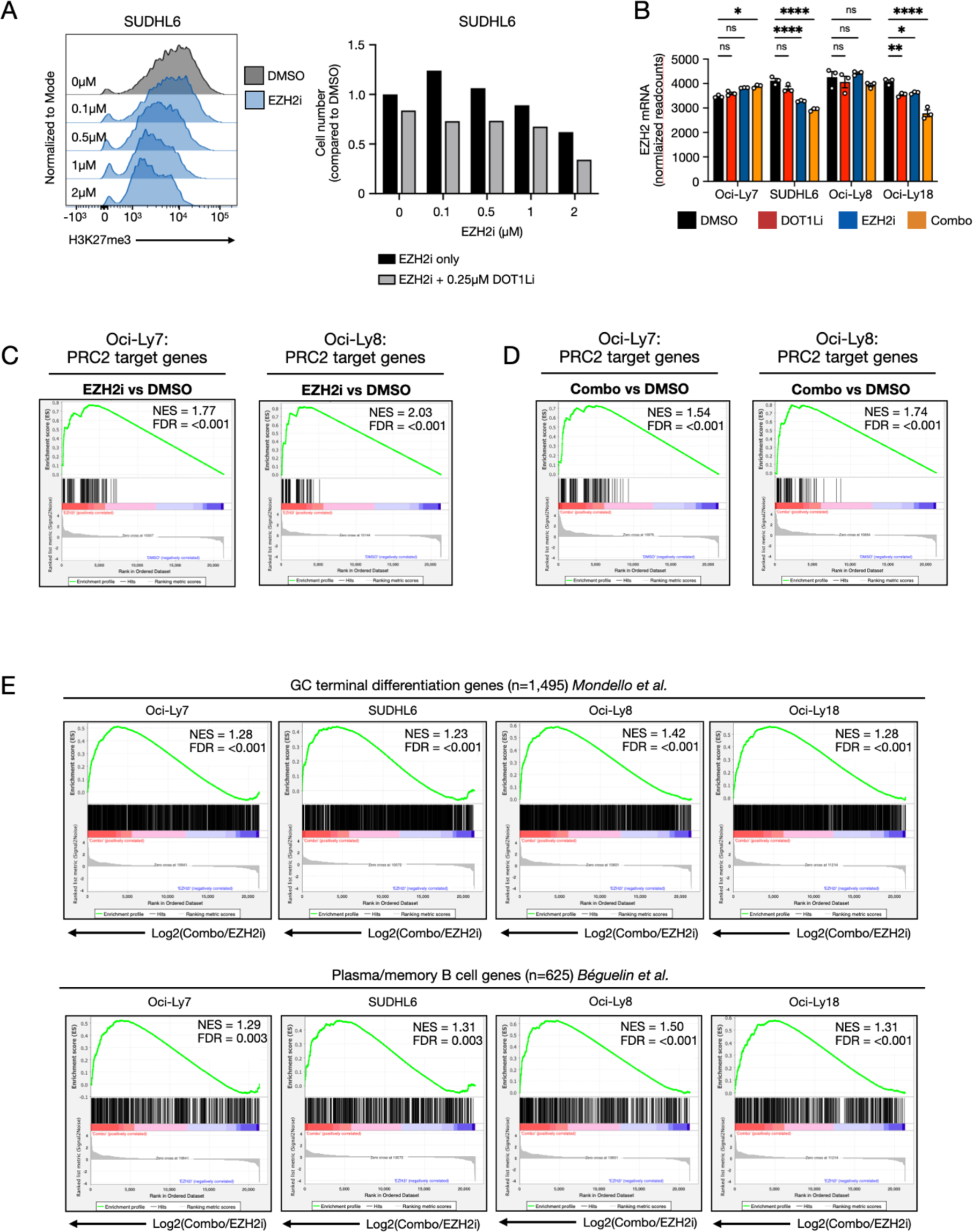
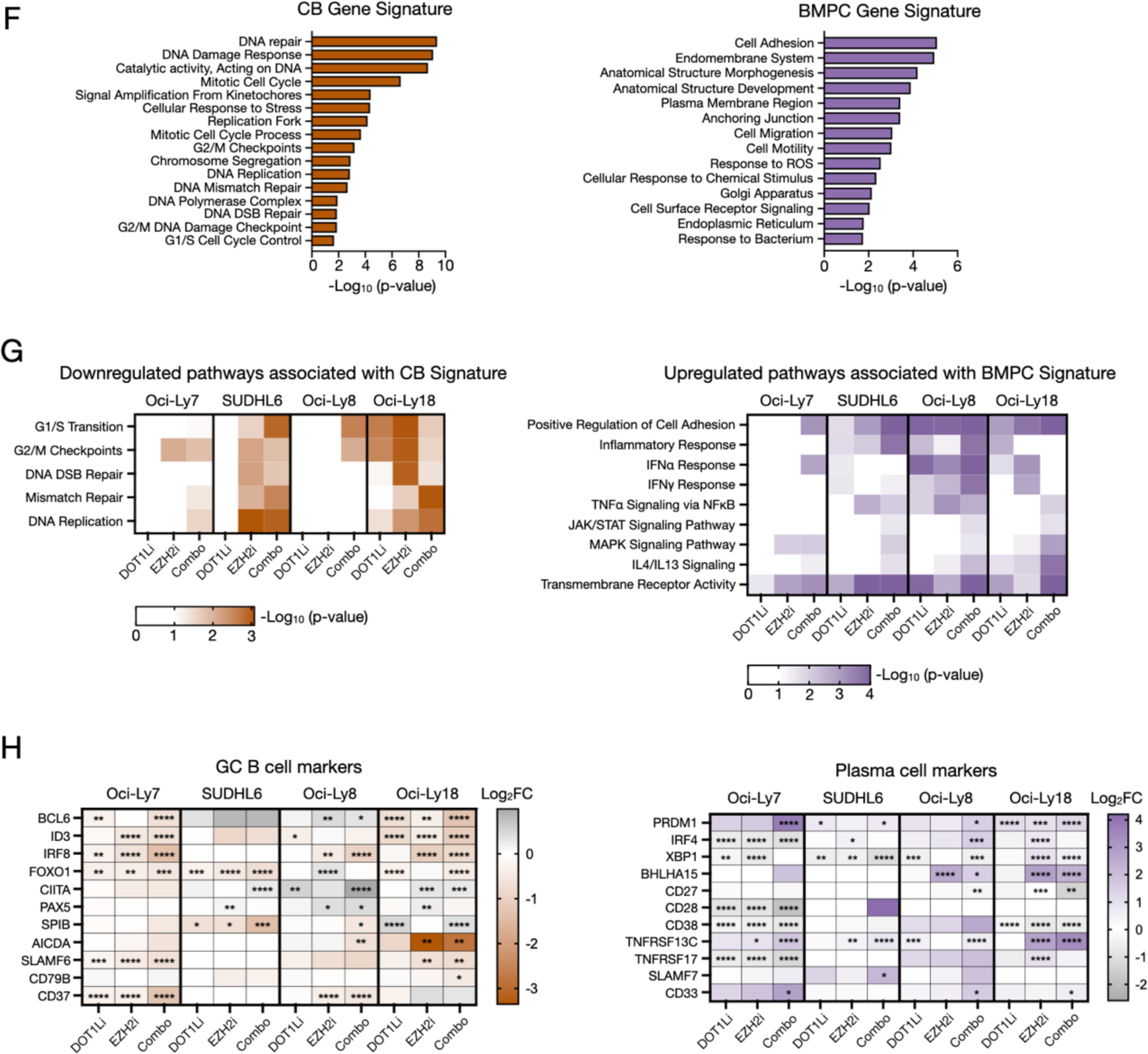
Combination treatment-induced derepression of PRC2 targets acts as driver for GCB-DLBCL to acquire a plasma cell-like state. (A) Representative histograms show the relative intracellular density of H3K27me3 after six days of treating SUDHL6 cells with different concentrations of EZH2i (left). The bar graph represents the effect on the number of cells upon titration of EZH2i alone or in combination with 0.25µM DOT1Li in SUDHL6 cells (right) (n=1). The DOT1Li concentration was chosen to maximally reduce H3K79me2 levels. (B) Normalized mRNA read counts of EZH2 acquired by RNA sequencing in four independent GCB-DLBCL cell lines treated with DMSO, DOT1Li alone, EZH2i alone, or combination of both treatments. (C) Confirmation that generated PRC2 target genes are enriched upon EZH2i using GSEA. (D) Similar to EZH2i alone, the combination treatment significantly derepresses PRC2 target genes. (E) Gene set enrichment analysis (GSEA) of germinal center terminal differentiation signature by *Mondello et al.*^30^, and GSEA of plasma/memory B cell signature by *Béguelin et al.*^11^ were conducted in four independent GCB-DLBCL cell lines. The GSEA plot indicates that the enrichment of a terminally differentiated plasma cell signature by the combination treatment is superior when compared to EZH2i alone. (F) Pathways enriched in CB signature (left) reflect the pro-proliferative identity of GC B cells, while BMPC gene signatures (right) are enriched with morphogenic and migratory alterations. (G) Downregulated pathways associated with CB signatures (left) and upregulated pathways associated with BMPC gene signatures (right) in four GCB-DLBCL cell lines upon treatment. (H) Differential expression of GC B cell (left) and plasma cell markers (right). Cells were treated for six days with the following concentrations of DOT1L1: 100nM for Oci-Ly7 and Oci-Ly8; 250 nM for SUDHL6; and 2µM for Oci-Ly18, and the following concentrations of EZH2i: 1µM for Oci-Ly7; 5µM for Oci-Ly8 and Oci-Ly18; and 2µM for SUDHL6. DOT1Li and EZH2i concentrations are based on the doses wherein H3K79me2 and H3K27me3 are maximally reduced. P-values were calculated using two-way ANOVA. ****P-value < 0.0001, ***P-value < 0.001, **P-value < 0.01, *P-value < 0.05. “ns” indicates not significant. NES = Normalized Enrichment Score. The false discovery rate (FDR) was calculated using GSEA. ***q-value < 0.001, ** q-value <0.01.

**Supplemental Figure 4.**
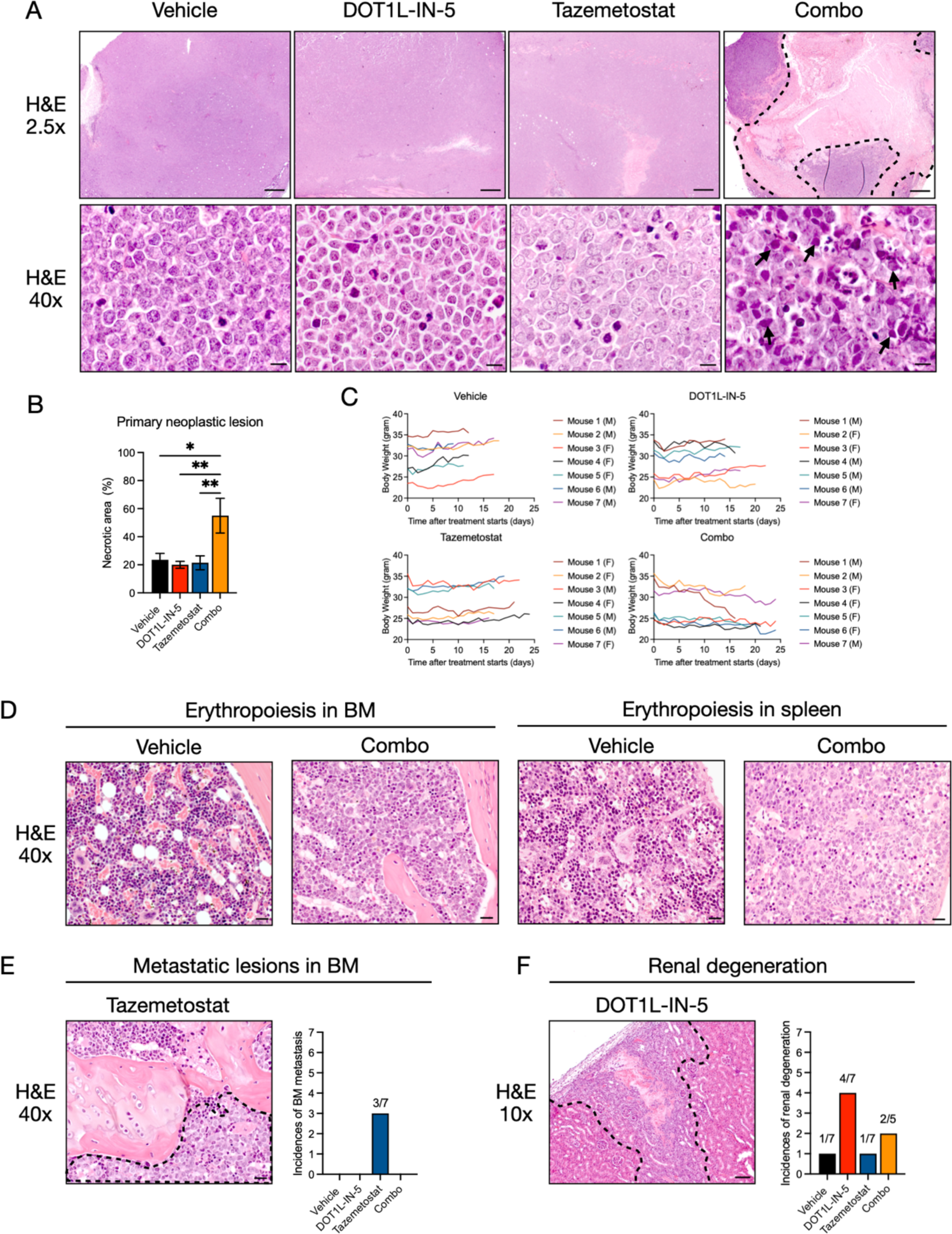
Combined inhibition of DOT1L and EZH2 induces apoptosis and necrosis *in vivo* accompanied by mild side-effects (A) Representative microphotograph images of hematoxylin and eosin (H&E) staining from Oci-Ly7 xenografts show an increase in apoptosis (arrow) and necrotic areas (dark dashed line) upon combination treatment. (B) Semi-quantitative analysis of necrotic areas by each treatment. (C) The body weight of all mice during the duration of the treatment with vehicle, DOT1L-IN-5, tazemetostat, and combination. Graphs indicates no effect of treatment on animal well-being. (D) Representative microphotograph images of H&E staining showing the depletion of erythropoietic cells (dark blue) in bone marrow and spleen upon combination treatment compared to vehicle. (E) Representative histopathological microphotograph images indicating the increase of metastatic lesions in bone marrow (BM) upon tazemetostat treatment (3 out of 7 mice) within the dark dashed line. (F) H&E staining of the kidney indicates nephrotoxicity in DOT1L-IN-5 (4 out of 7 mice) and combination treatment (2 out of 5 mice) within the dark dashed line. Magnification scale bar of microphotographs: 2.5x = 500µm, 10x = 100µm, 40x = 20µm. P-values were calculated using two-way ANOVA. **P-value < 0.01, *P-value < 0.05.

